# AAV2 Bypasses Direct Endosomal Escape by Using AAVR to Access the Trans-Golgi Network en Route to the Nucleus

**DOI:** 10.1101/2025.11.22.689972

**Authors:** Marti Cabanes-Creus, Sophia H.Y. Liao, Marta Pardo-Piñón, Adriana L. Rojas, Joseph Kelich, Jackson Coyne, Maddison Knight, Deborah Nazareth, Ramon Roca-Pinilla, Sujata Senapati, Matthew C. Walsh, Pedro J Cejas, Sean M Armour, Aitor Hierro, Leszek Lisowski

**Author notes:** Now at Pattern Biosciences, South San Francisco, CA, USA.

## Abstract

Vectors based on the adeno-associated virus are widely used as delivery methods in gene therapy applications, yet understanding of the mechanisms governing its intracellular trafficking remains incomplete. Traditional models suggest that AAV escapes from endosomes via membrane disruption, but direct evidence for this process are lacking. Here, we show that AAVR, the essential AAV cell entry receptor, functions as a bona fide retromer cargo. Using *in vitro* reconstitution assays, we demonstrate that AAVR’s cytosolic tail is sufficient to engage the SNX3–retromer complex and drive membrane tubulation, a hallmark of retrograde trafficking. In AAVR-knockout HuH-7 cells, AAV2 particles are internalized but fail to reach the *trans*-Golgi network (TGN) and support transgene expression. Galectin-8 recruitment assays reveal no evidence of endosomal membrane rupture during productive transduction, distinguishing AAV2 from lytic vectors such as lipid nanoparticles. Moreover, VP1u-deficient and PLA2-mutant AAV2 capsids accumulate at the TGN, indicating that VP1u is dispensable for early trafficking but required for post-TGN progression toward productive transduction. These findings challenge the prevailing model of direct endosomal escape and position AAV transduction as a vesicle-guided, receptor-mediated process.

## Introduction

Adeno-associated viruses (AAVs) are non-pathogenic, non-enveloped parvoviruses prevalent throughout the human population[1]. Wild-type AAV consists of a 4.7kb single-stranded DNA genome encoding for multiple genes, including non-structural (rep), structural (cap), assembly activating (aap), and membrane-associated accessory (maap) proteins[2, 3]. The structural proteins encoded by *cap*, VP1, VP2, and VP3, are assembled stochastically[4] into an icosahedral capsid, which encapsulates the viral DNA genome flanked by two inverted terminal repeats (ITRs)[4, 5]. In protototypical AAV2, VP1 and VP2 differ from VP3 by N-terminal extensions of 138 and 65 additional amino acids, respectively[6].

Although vectors based on AAV are currently the leading candidates for viral vector-based gene therapy approaches, basic aspects of AAV cellular entry and intracellular trafficking are still poorly understood[7]. The efficiency of both wild-type AAV infection as well as AAV vector transduction are limited at the cellular level by three major barriers: (i) cellular uptake, (ii) intracellular trafficking, and (iii) nuclear entry. Thus, a detailed study of the AAV infection/transduction process is of key importance for developing gene therapy vectors with improved safety and efficiency[6].

For many years, it was assumed that the primary receptor for AAV2, the prototypical and most studied AAV serotype, was the membrane-associated heparan sulfate proteoglycan (HSPG)[8]. This confusion resulted from the lack of distinction in the historical literature between the attachment of the AAV to the cell and the subsequent cellular entry. We now understand that from the virus’ and vector’s perspective alike, the glycans should not be considered as receptors, but rather as cell-attachment factors, interactions with which may not be highly specific[7, 9]. More recently, Pillay et al. shed some light on the mechanisms of cellular attachment and entry by identifying the protein receptor required for a functional AAV infection, the AAV-Receptor (AAVR)[7]. In contrast to HSPGs, AAVR has properties of a classical proteinaceous entry receptor[10]. Taking advantage of an unbiased haploid screen, Pillay and colleagues identified an uncharacterized type-I transmembrane protein, KIAA0319L (thereafter named AAVR), as a protein capable of rapid endocytosis from the plasma membrane and trafficking to the *trans*-Golgi network (TGN)[7]. The authors readily changed the paradigm of the field by showing that 1) AAVR directly binds to AAV2 particles, 2) anti-AAVR antibodies block AAV2 infection, and 3) genetic ablation of AAVR renders a wide range of mammalian cell types highly resistant to AAV2 infection[7]. Further structural studies from independent investigators confirmed the initial hypothesis on the key role that AAVR plays in AAV infection and transduction. Since its discovery, it has been confirmed that AAVR is utilized not only by AAV2, but also by other serotypes, with many structures of AAV-AAVR complexes resolved for AAV variants beyond prototypical AAV2[11, 12]. Thus, the currently accepted model of AAV cellular entry is that the initial attachment of vectors to the cell membrane, an event that utilizes cell surface exposed molecules, such as HSPG, is followed by engagement of the protein AAV receptor, which then dictates functional entry into the cell via endocytosis[10].

Once inside the cell, passage through acidic endosomal compartments has been traditionally associated with exposure of the N-termini of VP1 and VP2[13]. For AAV2, the VP1/2 N-termini contain a phospholipase A2 (PLA2) domain that was historically thought to be essential for endosomal escape[14]. However, while PLA2-mediated physical escape from endosomes has been generally considered as a possibility for the vector reaching the cytoplasm, direct evidence for this mechanism remains limited. Notably, PLA2 activity is dependent on calcium[15], and calcium levels may not be sufficient to trigger this activity, weakening the endosomal escape hypothesis[15]. After endosomal trafficking, AAV particles typically accumulate perinuclearly, a hallmark feature of parvovirus infection[6]. In the current model, it remains controversial how AAV breaches the nuclear envelope—some evidence suggests nuclear pore complex (NPC)-dependent entry[16], whereas other studies favor an NPC-independent mechanism[17, 18].

For vectors that bind strongly to HSPG, such as natural AAV3b and bioengineered AAV-KP1[19], we recently observed cellular entry but not transgene expression in a HuH-7 cell line harboring a knock-out of the AAVR gene[19]. This was not in contradiction to published data, given that only functional tests, specifically luciferase expression assays, were performed in the initial studies[7].

Here, we undertook studies to try to better understand the involvement of AAVR in AAV transduction. In this manuscript, we focus on the first step of functional transduction, trafficking from the plasma membrane to the TGN, to dissect the respective roles of AAVR and the N-terminal extension of VP1 (VP1u). Prototypical AAV2, which also binds strongly to HSPG[20], is an excellent tool because it can enter cells even in the absence of AAVR. Yet, consistent with previously published studies[7], functional transduction fails without AAVR involvement.

The data presented in this manuscript suggest that AAVR is specifically required for guiding AAV2 from the plasma membrane to the TGN, whereas VP1u, although essential at later steps, is not critical for this initial trafficking event. By combining immunofluorescence approaches with transmission electron microscopy, we provide evidence that the block to productive transduction observed in AAVR-KO cells occurs early in the intracellular pathway, caused by preventing AAV2 from reaching the TGN. Our results refine the current AAV infection paradigm by emphasizing the critical distinction between virus attachment via cell-surface glycans and the receptor-mediated trafficking to the TGN that is needed for successful transduction.

## Results

### AAVR is mainly localized at the *trans*-Golgi network at steady state, but can also be detected on the plasma membrane of non-permeabilized HuH-7 cells

To study the possible role of AAVR in AAV transduction, we took advantage of a previously described HuH-7 cell line harbouring a knock-out of the AAVR gene[19]. In the initial studies, we further characterized this cell line using immunofluorescence. In contrast to naïve HuH-7 cells, and as expected, we did not detect expression of AAVR in the knock-out cell line (**Fig. 1A-B**). As reported for HAP-1 cells[7], in naïve HuH-7 cells, we detected a strong association of AAVR with the *cis*-medial Golgi marker (giantin) (**Fig. 1A**), and a complete co-localization with the *trans*-Golgi network marker TGN46 (**Fig. 1C-D**). To investigate whether this intracellular localization was specific to HuH-7 cells, we also characterized AAVR expression pattern in primary human hepatocytes engrafted in a Fah^−/−^Rag2^−/−^Il2rg^−/−^ (FRG) mouse[21] (**Fig. 1E**) as well as in primary human hepatocytes in human liver sections (**Fig. 1F**). In both cases, at steady state, AAVR appeared to be mainly localized intracellularly. This is consistent with findings described in the original AAVR manuscript, where AAVR was only observed on the cell membrane when cells were maintained at 4°C[7].

**Figure 1.**
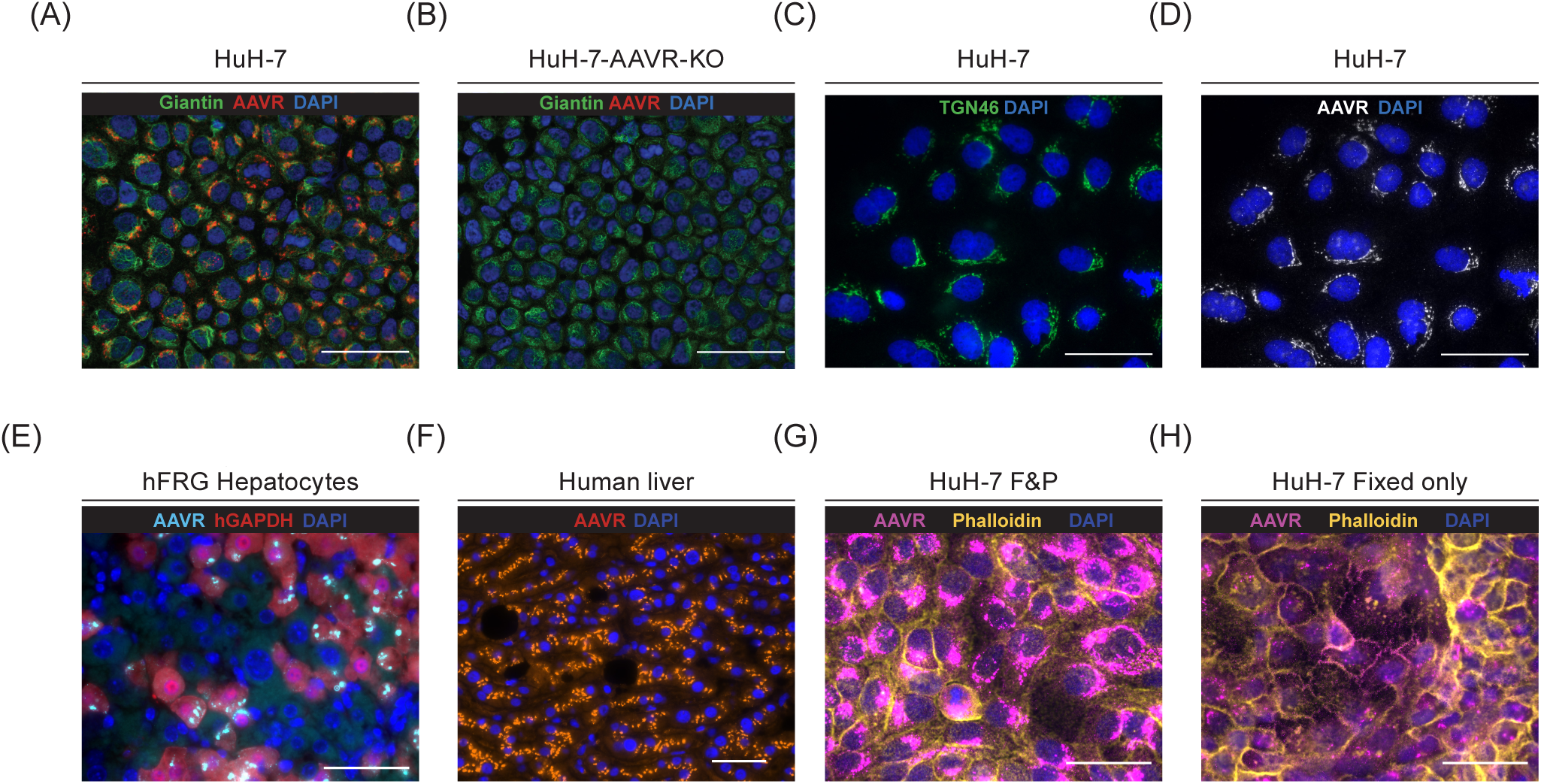
AAVR is mainly localized at the trans-Golgi network at steady state, but can be observed on the plasma membrane of non-permeabilized HuH-7 cells. **A–B)** Immunofluorescence images of naïve HuH-7 cells **(A)** and AAVR-knockout HuH-7 cells **(B)** showing colocalization of AAVR (red) with the Golgi markers Giantin (green), and DAPI (blue). AAVR is absent in knockout cells. **C–D)** Naïve HuH-7 cells stained for the trans-Golgi marker TGN46 (**C**, green) and AAVR (**D**, white), with DAPI in blue, confirming predominant intracellular localization of AAVR overlapping with TGN46. **E)** Humanized FRG liver sections showing human hepatocytes positive for intracellular AAVR (cyan) and human GAPDH (red). DAPI in blue. **F)** Human liver section stained for AAVR (orange) and DAPI (blue). **G–H)** HuH-7 cells stained for AAVR (magenta) and phalloidin staining of actin fibres (yellow), with DAPI in blue. Cells were either fixed and permeabilized **(G)** or only fixed without permeabilization **(H)**, showing that a subpopulation of AAVR is detectable at the plasma membrane under non-permeabilized conditions at 6 hours post-transduction. Scale bars: 50 µm.

Given that we routinely fix and permeabilize samples before antibody staining (**Materials and Methods**), we next investigated whether cell permeabilization affected our ability to visualize AAVR on the plasma membrane of naïve HuH-7 cells. In contrast to the mainly intracellular AAVR detected following fixing and permeabilization (**Fig. 1G**), we could readily detect AAVR on the plasma membrane, colocalizing with phalloidin staining of actin fibres, when the permeabilization step was omitted (**Fig. 1H**). These data indicate that an equilibrium exists between intracellular and extracellular AAVR, with a minor but detectable amount of AAVR present on the plasma membrane.

### AAVR is required for functional transduction, but not for AAV2 vector uptake, in HuH-7 cells

Next, we investigated the ability of AAV2 to transduce naïve and AAVR knock-out HuH-7 cells. To do so, we packaged a previously reported AAV expression cassette encoding eGFP under the control of liver-specific promoter (ss-LSP1-eGFP-WPRE-BGHpA)[22] into AAV2. To enable tracking vector particles intracellularly using immunofluorescence, we utilised the A20 antibody[23], which recognizes a conformational epitope present only in intact fully assembled AAV2 particles[24].

As observed in the time course experiment presented in **Fig. 2A-B**, we could detect intracellular AAV2 particles in both cell lines, regardless of the presence or absence of AAVR. The A20 signal peaked at 48 hours post-transduction and appeared lower at 72 hours, most likely due to proteasomal degradation. The most striking difference between both cell lines was the complete absence of eGFP signal in AAVR knockout cells (**Fig. 2B**). Given that we detected a strong cytoplasmic signal for intact AAV2 particles in the AAVR-KO cell line, we hypothesized that the lack of AAVR prevented AAV2 from trafficking to the TGN, a critical step *en route* to the nucleus.

**Figure 2.**
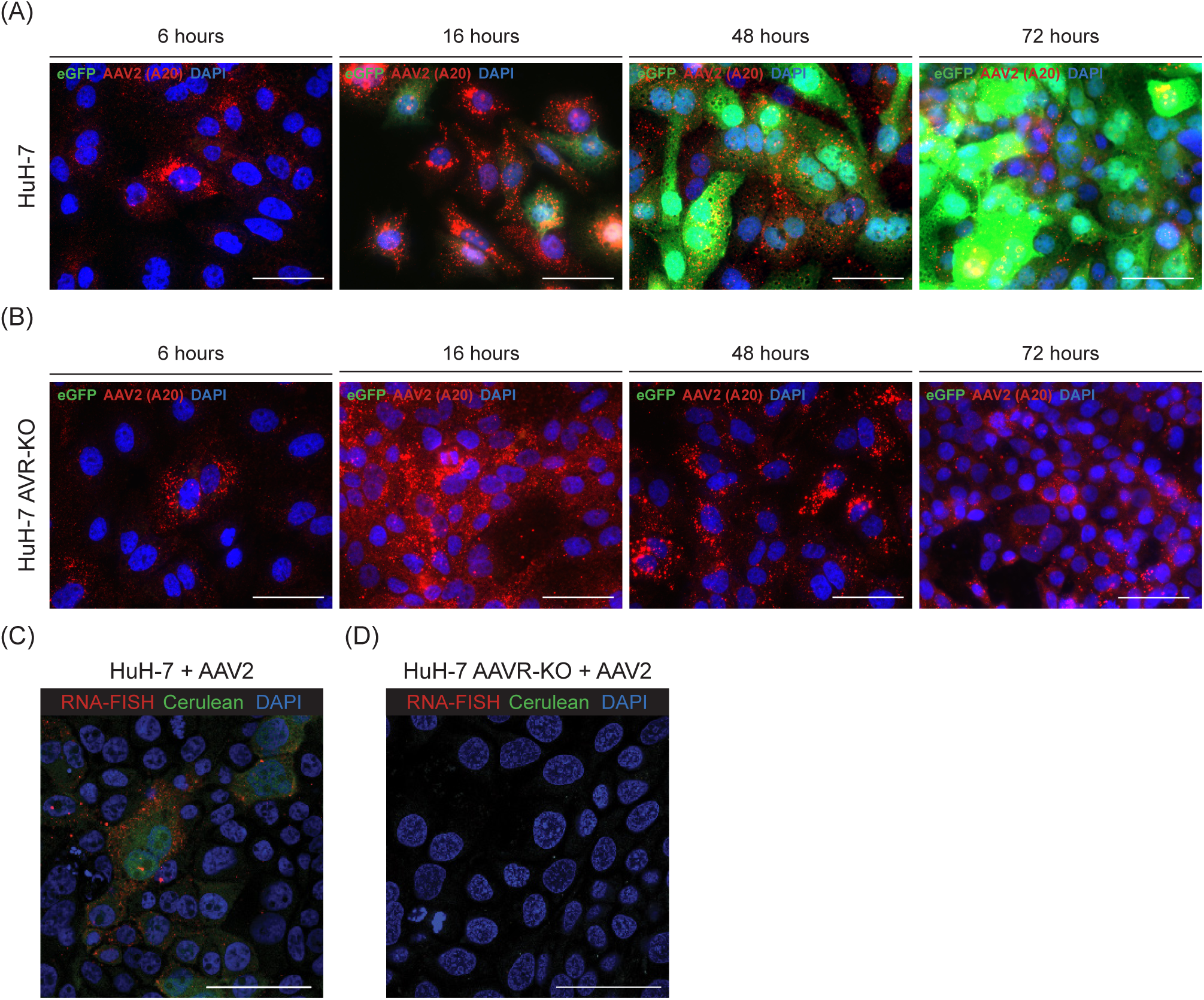
AAVR is required for a complete functional transduction but not for AAV2 vector uptake in HuH-7 cells. **A–B)** Time course of AAV2 vector uptake and eGFP transgene expression in naïve HuH-7 cells (**A**) and AAVR-knockout HuH-7 cells (**B**) at 6, 16, 48, and 72 hours post-transduction. AAV2 particles were detected using the A20 conformational antibody (red), eGFP expression marks successful transduction (green), and nuclei were stained with DAPI (blue). While AAV2 is internalized in both cell types, robust eGFP expression is observed only in AAVR-positive HuH-7 cells. **C–D)** RNA fluorescence in situ hybridization (RNA-FISH) detecting transgene mRNA (red) only in HuH-7 (**C**) but not in AAVR-knockout (**D**) cells transduced with an AAV2 vector encoding mCerulean. mCerulean protein (green) and DAPI (blue) were used to mark transduced cells and nuclei, respectively. Scale bars: 50 µm.

To determine whether this trafficking defect prevented the vector from reaching the nucleus and expressing the transgene, we transduced naïve and knockout cell lines with an AAV2 vector encoding mCerulean fluorescent reporter and assessed transgene expression at the mRNA level using high-resolution RNA fluorescence *in situ* hybridization (RNA-FISH). In contrast with the naïve cells (**Fig. 2C**), we could not detect transgene RNA in the AAVR knock-out HuH-7 cell line (**Fig. 2D**). Consistent with the absence of transgene mRNA, the mCerulean fluorescent reporter was also not detectable at the protein level.

These data indicate that AAVR is essential for intracellular trafficking events required for AAV transduction before nuclear entry. Given the established role of *trans*-Golgi network trafficking in productive AAV transduction[15], we next examined the intracellular route of AAV following cell entry, focusing on three key aspects: 1) the mechanism of AAVR-dependent trafficking, 2) the possibility of direct endosomal escape, and 3) the specific stage within the trafficking pathway at which VP1u becomes functionally required.

### AAVR is a retromer cargo capable of mediating endosomal tubulation, supporting a mechanistically plausible route for AAV trafficking to the TGN

Previous studies demonstrated that trafficking to the *trans*-Golgi network is required for productive AAV transduction[15], although the specific role of AAVR in this process remains undefined. Notably, the original genome-wide screen that identified AAVR as an essential receptor for AAV transduction[7] also revealed multiple hits associated with Golgi and TGN function, implicating this compartment as a critical component of the AAV infection pathway. Based on these observations, we hypothesized that AAVR may function as a cargo of the retromer complex, a multi-protein assembly that associates with the cytosolic surface of endosomes and mediates the recycling of membrane proteins, including receptors and transporters, to the TGN via tubular-vesicular carriers[25].

To test this hypothesis, we first investigated the role of AAVR’s cytosolic tail, as cytosolic domains of many transmembrane proteins often harbor trafficking motifs that are recognized by the cellular sorting machinery, thereby regulating endocytosis and intracellular routing of the receptor[25]. We expressed a truncated version of AAVR lacking its cytosolic tail in AAVR-knockout cells. Unlike the wild-type protein, which localized predominantly to the TGN, the truncated variant exhibited a dispersed cytoplasmic distribution (**Fig. 3A-B**), suggesting that the cytosolic tail is required for proper subcellular localization. These results are consistent with a model in which AAVR relies on specific cytosolic sorting signals to engage intracellular trafficking pathways. Notably, near the transmembrane domain of AAVR, there is a putative retromer-SNX3 binding motif (Ωx[L/M/V])[25], where Ω denotes a bulky hydrophobic residue, corresponding in this case to the sequence YKI (residues 968-970). Structural predictions using AlphaFold3[26] further suggest that this region adopts a conformation analogous to that of the DMT1-II endosomal sorting signal when bound to the retromer-SNX3 complex[25] (**Fig. 3C–F**).

**Figure 3.**
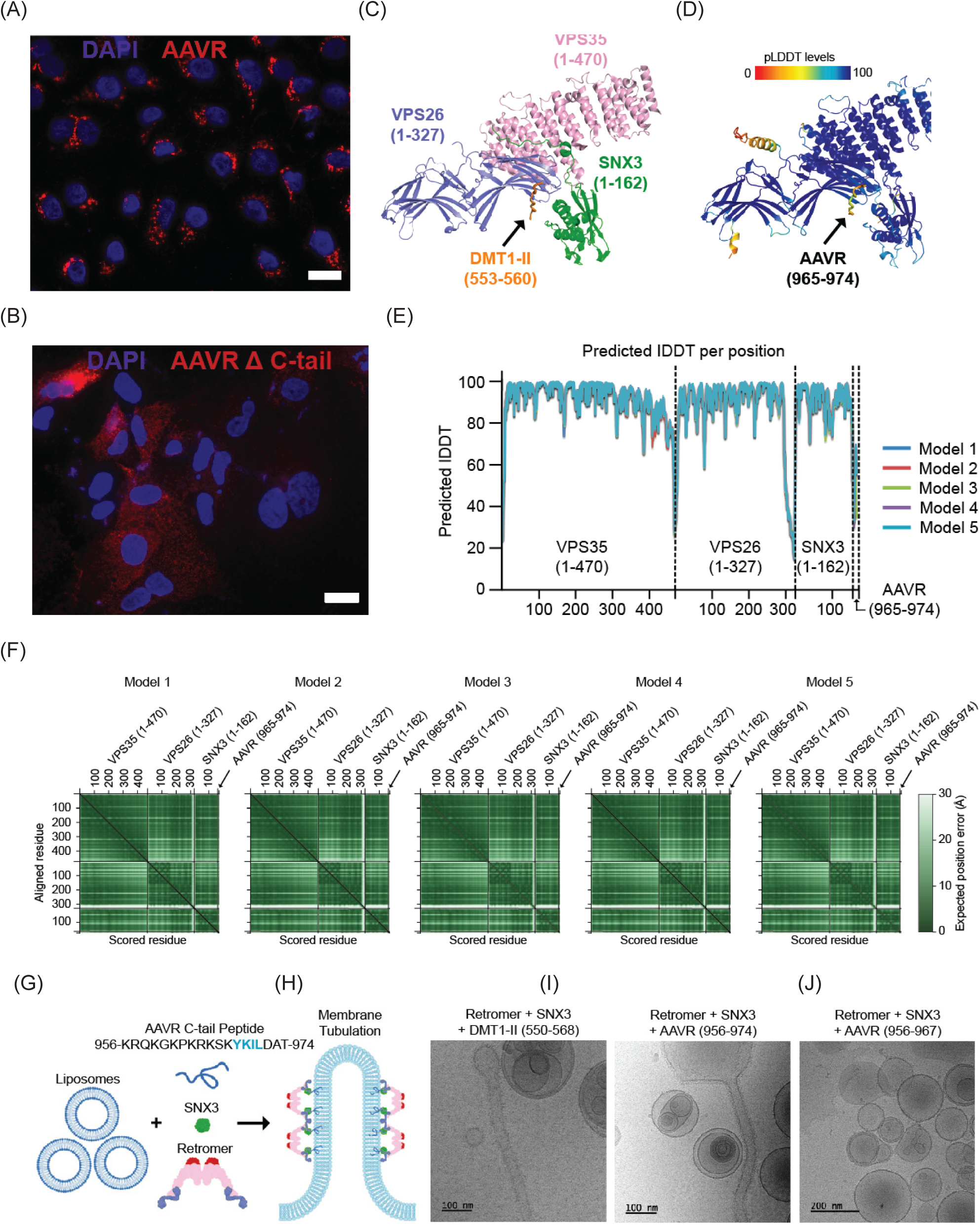
AAVR is a SNX3–retromer cargo that could mediate AAV trafficking to the trans-Golgi network via endosomal tubulation. **A–B)** Confocal immunofluorescence images of AAVR localization in AAVR-knockout HuH-7 cells reconstituted with either wild-type AAVR (**A**) or a truncated AAVR lacking the cytosolic tail (ΔC-tail; **B**). Wild-type AAVR shows punctate intracellular localization consistent with TGN localization, while the ΔC-tail variant displays diffuse cytoplasmic staining. Nuclei are stained with DAPI (blue); AAVR is shown in red. **C**) Crystal structure of the DMT1-II cargo peptide bound to the retromer-SNX3 complex (PDB: 5F0L[25]), shown for comparison with **D**, the AlphaFold3-predicted model of the AAVR cytosolic tail peptide (residues 956-974) bound to retromer–SNX3. In **D**, the model is colored by per-residue confidence score (pLDDT; scale 0-100). **E-F**) Quality estimation metrics for the top five AlphaFold3 models. **E**, per-residue pLDDT confidence plots; **F**, Predicted Alignment Error (PAE) plots for VPS35 (residues 1-470), SNX3 (1-327), and AAVR (956-974) complexes. **G)** Schematic of the *in vitro* reconstitution assay used to test AAVR as a retromer cargo. A peptide derived from the AAVR cytosolic tail containing a candidate retromer-binding motif was incubated with liposomes, SNX3, and retromer complex components. Successful recruitment of retromer components to the membrane induces curvature and tubule formation (∼35 nm diameter). **H–J)** Transmission electron microscopy (TEM) images of liposomes incubated with retromer and SNX3 in the presence of a known cargo peptide from DMT1-II (**H**), the AAVR cytosolic tail peptide (residues 956-974) (**I**), or a mutant peptide lacking the predicted sorting signal (residues 956-967) (J). Both DMT1-II and AAVR peptides induce the formation of narrow membrane tubules, consistent with retromer-driven membrane tubulation. Scale bars: 20 µm (A–B); 100 nm (H-I); 200 nm (J).

To directly assess whether AAVR functions as a retromer cargo, we employed an *in vitro* reconstitution system using liposomes and purified retromer components, based on previously established protocols designed to study the structure of retromer–SNX3 assemblies on membranes[27, 28] (**Fig. G**). We adapted the assay by systematically limiting the number of components to evaluate their capacity to induce liposome tubulation. Consistent with a cargo-dependent mechanism, only the inclusion of the known DMT1-II cargo peptide enabled membrane tubulation, while all other combinations of individual components or pairwise mixtures failed to induce remodeling. Similarly, a peptide derived from the cytosolic tail of AAVR (residues 956–974), which includes the predicted retromer recognition motif, also triggered tubule formation (**Fig. H, I,** and **Supplementary Fig. 1**). In contrast, deletion of the motif within this sequence (AAVR 956–967) abolished tubulation (**Fig. J**), highlighting the functional importance of this sorting signal for retromer-mediated membrane remodeling. The tubules formed in these reactions exhibited an average diameter of ∼35 nm, consistent with the physical dimensions required for AAV particle trafficking.

These findings demonstrate that the cytosolic tail of AAVR contains a sorting motif capable of engaging the retromer–SNX3 complex and initiating membrane remodeling *in vitro*. While this *in vitro* system does not confirm retromer-mediated AAV trafficking to the TGN in cells, it supports a mechanistically plausible route aligned with known retromer functions and the spatial constraints of AAV transport.

### AAV2 does not show evidence of direct endosomal escape during productive transduction

To assess whether AAV2 escapes the endosomal system via membrane disruption or through vesicular trafficking, we employed a Galectin-8 (Gal8)-based assay to monitor endosomal integrity. Gal8 is a cytosolic lectin that binds to β-galactoside glycans, which are normally restricted to the luminal face of endosomal membranes. Upon endosomal rupture, these glycans become exposed to the cytosol, leading to Gal8 recruitment and the formation of fluorescent puncta when visualized in cells expressing Gal8-GFP[29, 30]. This assay has been validated as a sensitive method to detect endosomal disruption caused by viruses, nanoparticles, or other membrane-lytic agents[31].

We transduced eGFP-Gal8-expressing HepG2 cells with AAV2 vectors encoding mScarlet at multiplicities of transduction (MOTs) ranging from 10³ to 10⁵ vector genomes per cell and monitored Gal8 puncta formation over time using live-cell fluorescence microscopy. Across all doses tested, AAV2 failed to induce a detectable increase in Gal8 puncta compared to untreated controls (**Fig. 4A-C**), indicating no measurable endosomal membrane disruption. In contrast, treatment with mRNA-loaded lipid nanoparticles (LNPs), which are known to escape endosomes via membrane lysis, resulted in robust Gal8 puncta formation, confirming the sensitivity of the assay (**Fig. 4A-C**).

**Figure 4.**
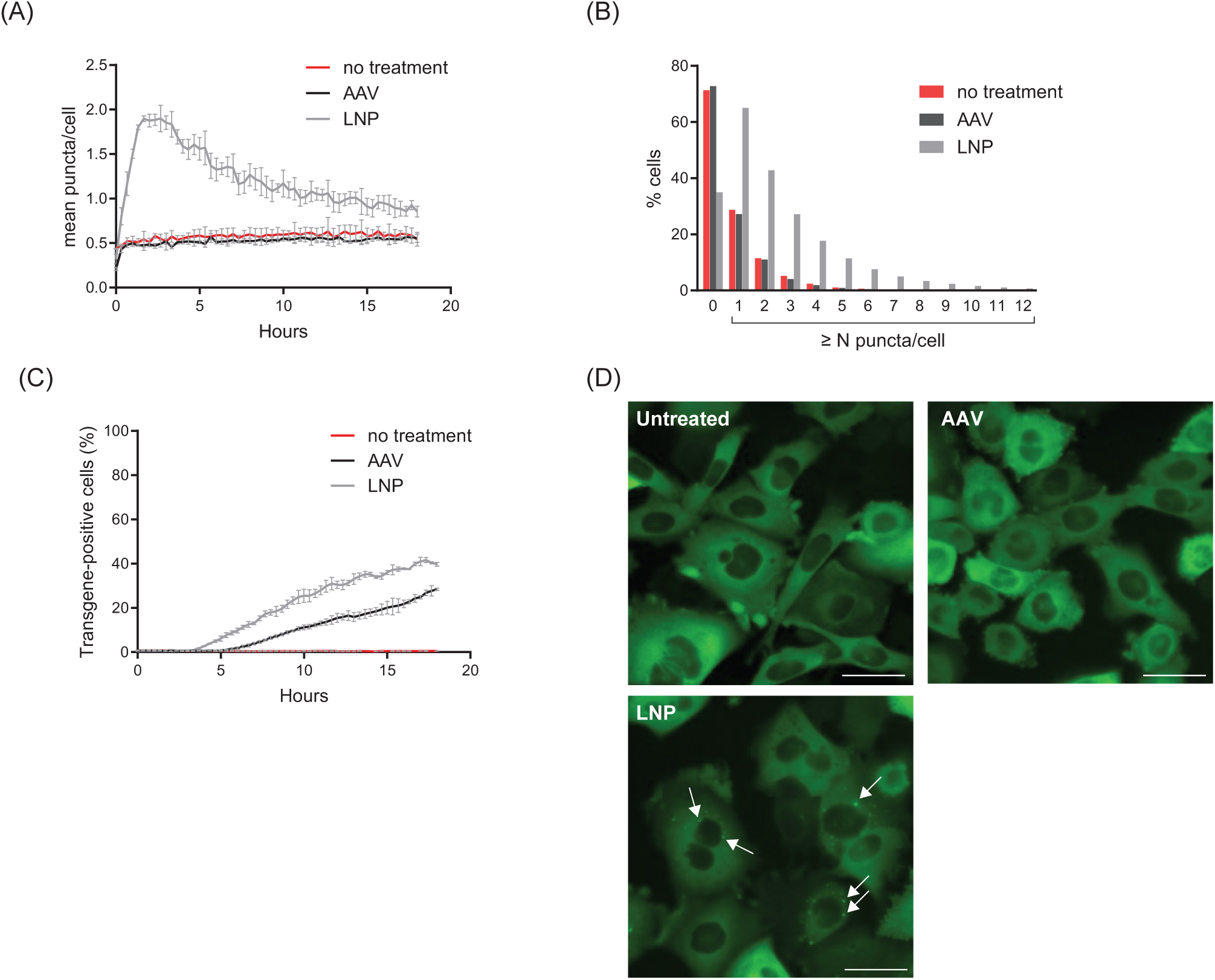
AAV2 does not induce Galectin-8 recruitment, indicating a lack of endosomal membrane disruption during productive transduction. **A)** Time course of mean Gal-8 puncta per cell quantified from live-cell timelapse imaging of eGFP-Gal 8 signal from eGFP-Gal8-HepG2 cells. Data are shown as mean of 3 treatment groups per condition. 11 ng per well dosing was used for LNP while 10⁵ MOT was used for AAV dosing. **B)** Cumulative frequency distribution representing Gal-8 puncta per cell at 2.3 hours post-treatment. N = 4748 for the no treatment group, N = 2829 for the AAV group, and N = 4693 for the LNP-treated group. **C)** Representative microscope images taken at 2.3 hours post-treatment showing eGFP Gal-8 distribution in mRNA-LNP-treated, AAV-treated, or untreated (no treatment). **D)** Time course is shown for the percentage of transgene-positive cells quantified from mCherry or mScarlett fluorescence for mRNA-LNP or AAV, respectively. Data are shown as the mean of n=3 treatment groups per condition.

Despite the absence of Gal8 recruitment, AAV2-mediated transgene expression was detected over time, with a progressive increase in the percentage of transgene-positive cells up to 18 hours post-transduction (**Fig. 4D**), indicating successful nuclear delivery and productive transduction. These findings indicate that productive AAV2 transduction does not involve substantial endosomal membrane disruption. Instead, they support a model in which AAV2 traffics through the endosomal system to the *trans*-Golgi network, consistent with the AAVR-and retromer-dependent pathway described above. This mechanism differs from the cytosolic escape strategy characteristic of other gene delivery platforms, such as LNPs.

### VP1u and its PLA2 activity are required for post-TGN trafficking during productive AAV2 infection

The absence of transgene expression when using wild-type AAV2 capsids in AAVR-knockout cells (**Fig. 2A-B**) closely resembles the phenotype observed with AAV2 capsids lacking the VP1 unique region (VP1u), which are capable of cellular entry but fail to mediate productive transduction[6]. To investigate the relationship between AAVR-mediated trafficking and VP1u function, we packaged a liver-specific eGFP expression cassette (ss-LSP1-eGFP-WPRE-BGHpA) into either wild-type AAV2 capsids or VP1-deficient (VP2/VP3-only) AAV2 particles. While AAV2 achieved robust transduction, VP1-deficient vectors did not lead to a detectable transgene expression, despite clear perinuclear accumulation (**Fig. 5A-B**).

**Figure 5.**
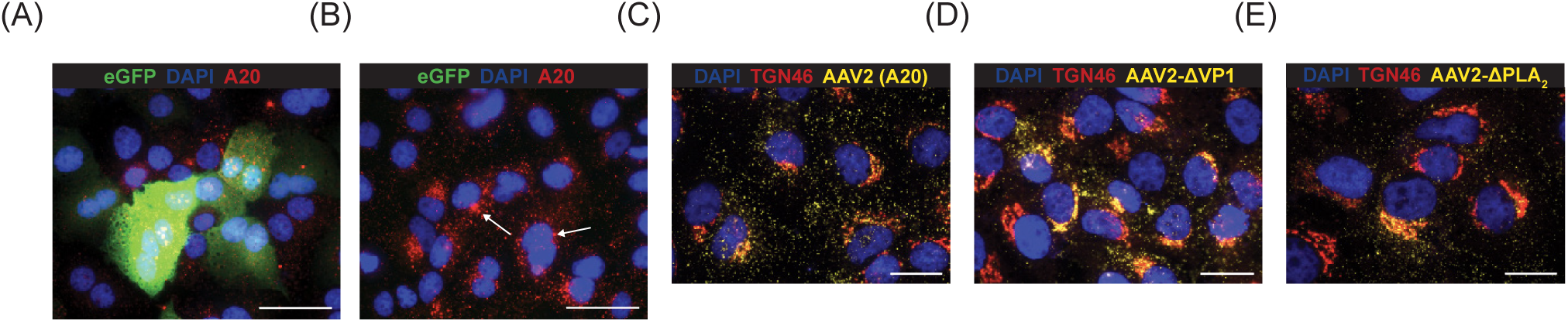
VP1u-deficient particles reach but accumulate at the trans-Golgi network, suggesting a post-TGN essential role. **A–B)** AAV2 **(A)** or ΔVP1-AAV2 **(B)** capsid uptake and eGFP transgene expression in HuH-7 cells at 48 hours post-transduction. AAV2 particles were detected using the A20 conformational antibody (red), and eGFP expression (green) marks successful transduction. DAPI (blue) marks nuclei. While wild-type AAV2 shows robust eGFP expression at 48 hours post-transduction, ΔVP1-AAV2 particles are internalized but fail to support transgene expression, accumulating in perinuclear puncta (white arrows). **C–E)** Immunofluorence images showing colocalization of internalized capsids (A20) with the trans-Golgi marker TGN46 (red) at 24 hours post-transduction in HuH-7 cells transduced with wild-type AAV2 (**C**, yellow), ΔVP1-AAV2 (**D**, yellow), or PLA2-deficient AAV2 (**E**, yellow). DAPI (blue) labels nuclei. Both ΔVP1 and PLA2 mutant particles show strong overlap with TGN46, indicating successful trafficking to the TGN but failure to proceed beyond it. Scale bars: 50 µm (**A–B**), 20 µm (**C–E)**.

To further characterize this defect, we analyzed the subcellular localization of wild-type AAV2, VP1-deficient AAV2, and a PLA2 catalytic mutant (H75A/D76N)[6] in HuH-7 cells. Both the VP1-deficient and PLA2-mutant particles showed enhanced colocalization with *trans*-Golgi network (TGN) markers compared to AAV2 at 24 hours post-transduction, indicating that these variants can reach the TGN but are unable to proceed beyond it (**Fig. 5C-E**). These findings support a model in which AAVR mediates trafficking of AAV to the TGN independently of VP1u, but VP1u and its PLA2 enzymatic activity are required for a downstream post-TGN step in the infectious pathway.

## Discussion

Our findings support a model in which AAV traffics to the *trans*-Golgi network (TGN) via direct interaction with AAVR, which itself functions as a cargo of the retromer complex. This challenges traditional views of AAV intracellular transport and prompts a re-evaluation of the long-standing concept of “endosomal escape.” To critically assess this idea, it is important to revisit its origins in the AAV field and examine the experimental evidence that has been used to support it.

The concept of “endosomal escape” in the AAV field is widely accepted despite a lack of direct experimental evidence supporting this model. Traditionally, “endosomal escape” has been interpreted as the direct penetration of the endosomal membrane by the viral particle and AAV release into the cytosol, similar to mechanisms described for adenovirus[32]. However, this interpretation may be over simplified or narrow. In the case of AAV, alternative trafficking mechanisms, such as the regulated transport of viral particles from endosomes to the *trans*-Golgi network (TGN) via host sorting machinery, may fulfill the functional equivalent of “endosomal escape.” This broader definition is supported by our findings and previous work[33] demonstrating that AAV traffics to the TGN, likely through hijacking the AAVR–retromer pathway.

More than a decade ago, Nonnenmacher and Weber pioneered the concept that productive AAV transduction requires passage through the TGN and identified Golgi-dependent trafficking as a critical step in the infectious pathway of AAV2[33, 34]. While their work established the importance of retrograde transport, the molecular link between the endosomal system and TGN targeting remained unclear. We propose that AAVR, as a retromer cargo, represents the missing mechanistic connection between membrane entry and TGN-directed trafficking in the AAV lifecycle. Notably, the genome-wide haploid screen conducted by Pillay et al.[7] identified multiple components of the retromer complex as essential for AAV2 infection, directly supporting a model in which AAV exploits host sorting machinery to reach the TGN. In contrast, the earlier study by Nonnenmacher et al. emphasized clathrin-independent entry via the CLIC/GEEC pathway and also observed TGN trafficking, but did not report a strict dependence on retromer components[34]. These seemingly divergent conclusions may reflect differences in experimental systems, where Pillay et al. employed HAP1 cells (haploid cells enabling complete gene knockouts), providing definitive disruption and lack of protein expression, whereas Nonnenmacher et al. used transient knockdowns and pharmacological inhibitors, which may have led to incomplete protein removal. Alternatively, these findings may not be mutually exclusive: AAV may exploit multiple, parallel pathways to access the TGN, with one involving AAVR-retromer-mediated sorting and another proceeding through CLIC/GEEC endocytosis. Such dual trafficking routes could reflect cellular redundancy or capsid serotype-specific preferences and underscore the complexity of AAV intracellular transport.

The prevailing notion of direct endosomal escape in the AAV field may have been influenced by studies on canine parvovirus (CPV), a related member of the Parvoviridae family. CPV was shown to induce pore formation during endocytic entry via the phospholipase A2 domain located in the N-terminus of VP1[35]. This PLA2 activity, triggered by acidic pH, was proposed to mediate membrane permeabilization. However, CPV-induced pores only allowed the cytosolic translocation of dextrans up to 3 kDa, while larger 10 kDa dextrans remained trapped within endosomes. Moreover, coendocytosed α-sarcin, a larger protein, was also retained in endosomes. These findings suggest that while CPV can alter endosomal membrane permeability[35], it does not cause vesicle rupture or enable passage of particles comparable in size to intact AAV (∼25 nm).

In this work, we show that AAVR functions as a retromer cargo and is sufficient to drive membrane tubulation *in vitro* through SNX3–retromer–dependent remodeling, a process consistent with retrograde trafficking to the TGN (**Fig. 3**). This potential trafficking mechanism aligns with prior electron microscopy observations of AAV5 in tubular endosomal structures[36]. Additionally, we demonstrate that AAV2 transduction does not induce detectable endosomal membrane rupture, as measured by a Galectin-8 reporter assay, even under high vector loads that nonetheless result in productive transgene expression (**Fig. 4**). These findings collectively support a model in which AAV exploits regulated vesicular pathways, rather than membrane lysis, to navigate intracellular barriers and reach the nucleus.

Importantly, our data do not provide additional mechanistic insight into how VP1u mediates AAV trafficking. However, they offer an important spatial and temporal clue: VP1u-lacking or PLA2-deficient AAV virions successfully traffic to the *trans*-Golgi network (TGN) but fail to progress further, instead accumulating at this compartment (**Fig. 5C-E**). This observation suggests that the role of VP1u becomes critical specifically at or beyond the TGN. In line with this model, GPR108, a highly conserved AAV entry factor localized in the Golgi, has been shown to interact functionally with the VP1u domain, and its absence results in impaired nuclear import despite intact endosomal uptake[37]. These findings together raise the possibility that VP1u becomes functionally relevant upon retrograde trafficking to the TGN. Following this checkpoint, it remains to be elucidated whether AAV proceeds to the nucleus via continued vesicular transport through the endoplasmic reticulum (ER) or whether the viral particle physically enters the cytosol before nuclear import.

A second central question in AAV biology, with profound implications for both basic virology and gene therapy applications, is where AAV first engages its essential receptor AAVR. While the requirement for AAVR in productive AAV infection/transduction is clear, the site of this critical interaction remains unclear: does AAV bind AAVR at the cell surface to initiate its intracellular journey, or does this crucial interaction occur only after internalization? This seemingly straightforward question has proved surprisingly challenging to resolve, in part because, as mentioned before, AAV2 can enter cells through multiple pathways, not all of which lead to successful infection or transduction. Understanding the distinction between productive and non-productive entry pathways has proved key to untangling this complex question.

Our findings highlight a critical distinction in AAV cellular entry pathways that help resolve apparent contradictions in the field: AAV2 can enter cells through both AAVR-dependent and AAVR-independent mechanisms, but only AAVR-mediated entry leads to successful transduction (**Fig. 2A-B**). The AAVR-independent entry, likely occurring through general cellular uptake mechanisms such as micropinocytosis, represents a “dead end” pathway that fails to establish productive transduction despite successful internalization. This distinction between functional and non-functional entry helps explain why early studies focusing solely on viral entry could be misleading. Several lines of evidence support this two-pathway model. The original work by Pillay et al. that led to the identification of AAVR showed that anti-AAVR antibodies efficiently blocked productive AAV2 infection when pre-incubated with cells, reducing functional transduction by over 10-fold[7]. This blocking effect suggests that AAV-AAVR interactions at the plasma membrane are crucial for initiating the productive entry pathway. Additionally, soluble AAVR protein could compete with cellular AAVR and neutralize AAV2 infection in a dose-dependent manner, further supporting the importance of these early interactions for successful transduction[7]. In a more recent *in vivo* study, vector DNA was detectable but RNA was markedly reduced to faint or undetectable levels in AAVR-knockout murine hepatocytes, supporting a non-productive entry pathway[38]. While the vector can still enter cells through AAVR-independent mechanisms, it fails to establish the proper trafficking route needed for successful gene expression.

This model also aligns with our immunofluorescence data showing AAVR’s distribution between the plasma membrane and *trans*-Golgi network. While AAVR is predominantly localized to the TGN at steady state, the detectable pool of AAVR on the plasma membrane may serve to engage AAV particles early and direct them into the productive trafficking pathway, distinguishing them from particles that enter through non-specific mechanisms (**Fig. 1G-H**). Also, recently, *in vivo* studies using the SELECTIV mouse system have provided additional compelling evidence for AAV-AAVR interaction occurring at the cell membrane[39]. Biodistribution studies using radiolabeled AAV9 demonstrate that mice expressing AAVR only in cardiac tissue show rapid and specific accumulation of viral particles in the heart within hours of systemic delivery[39]. This immediate tissue tropism strongly suggests AAVR functions as a cell surface receptor to capture and concentrate viral particles from circulation, as such rapid tissue-specific accumulation would not be expected if the critical interaction occurred only after internalization. These *in vivo* data reinforce the concept that AAVR functions as a gatekeeper at the cell surface, determining which cells can initiate productive entry.

More recently, Dhungel et al. identified carboxypeptidase D (CPD) as a host factor that supports the transduction of clade E AAVs (AAV8, AAVrh10, AAVhu37) in the absence of AAVR and serves as an essential entry factor for AAV11 and AAV12, which transduce cells independently of AAVR[40]. For AAV8, while CPD can substitute for AAVR when overexpressed, its loss alone does not impair AAV8 transduction. CPD is a Golgi-resident protein that cycles between the plasma membrane and trans-Golgi network, similar to AAVR, and can deliver AAV capsids to the same intracellular compartment required for productive infection. These findings are compelling, though we suggest caution in their broader interpretation. In virological convention, receptor nomenclature typically reflects an essential and conserved interaction that is functionally required across viral species or serotypes. Since CPD can substitute for AAVR in certain contexts, particularly for a subset of clade E capsids and primarily in AAVR-deficient settings, it may be more appropriate to describe it as a serotype-restricted entry factor rather than as a canonical “AAVR2”. Moreover, CPD likely did not evolve to interact with AAV, but rather fulfills its own defined cellular functions, making its engagement by specific AAVs a case of opportunistic rather than co-evolved usage. This distinction helps preserve conceptual clarity between essential and auxiliary host dependencies. Nevertheless, this discovery presents an exciting engineering opportunity: because CPD shares key localization features with AAVR, namely cycling between the plasma membrane and the TGN, it demonstrates that other host proteins in this trafficking pathway can substitute for AAVR if the capsid is appropriately tuned. We therefore propose that future capsid engineering efforts could deliberately reprogram receptor usage away from AAVR and toward alternative factors such as CPD, potentially enabling more tissue-restricted or cell-type-specific transduction.

While our mechanistic insights provide a refined model for AAV intracellular trafficking, it is important to note that all experiments in this study were performed with AAV2. The extent to which these findings generalize to other serotypes remains to be determined, as differences in receptor usage, glycan binding, and entry pathways can substantially alter intracellular routing. In particular, the degree of AAVR-independent, non-productive “dead entry” is almost certainly serotype dependent. Capsids with strong heparan sulfate proteoglycan (HSPG) affinity, such as AAV2, AAV3b, and AAV6, may be especially prone to uptake through non-productive routes that fail to engage AAVR. Notably, in non-human primates, AAV2 and AAV6 elicit markedly stronger ocular immune responses than non-HSPG-binding serotypes, consistent with enhanced diversion into degradative pathways that increase capsid presentation and immunogenicity[41]. We therefore hypothesize that the heightened immunogenicity often attributed to AAV2 *in vivo* may stem, at least in part, from a larger fraction of capsids being sequestered and degraded in these dead-end compartments. Taken together, this serotype-specific context should be considered when interpreting our results, which should be viewed as AAV2-specific while providing a conceptual framework for understanding core principles of AAV entry and trafficking.

In summary, our work reveals that AAV can enter cells through AAVR-independent pathways, but these routes lead to a dead end, failing to support productive infection. We propose a refined model for productive AAV trafficking (**Fig. 6**): following initial attachment to cell surface proteoglycans, AAV engages AAVR at the plasma membrane, initiating receptor-mediated endocytosis. We show that AAVR functions as a cargo of the retromer complex, and that AAV likely exploits retromer-mediated tubular transport to traffic from endosomes to the *trans*-Golgi network. Importantly, we find no evidence for direct endosomal membrane disruption by AAV, as measured by Galectin-8 recruitment, challenging the conventional model of endosomal escape. Our data demonstrate that VP1u and its PLA2 activity are dispensable for trafficking to the TGN, as VP1u-deficient and PLA2-mutant particles successfully reach this compartment but accumulate there without progressing further. This indicates that VP1u function becomes essential specifically for post-TGN trafficking steps required for productive transduction. This model highlights a previously underappreciated spatial checkpoint at the TGN that separates AAVR-dependent retrograde transport from VP1u-dependent downstream trafficking in the AAV transduction pathway.

**Figure 6.**
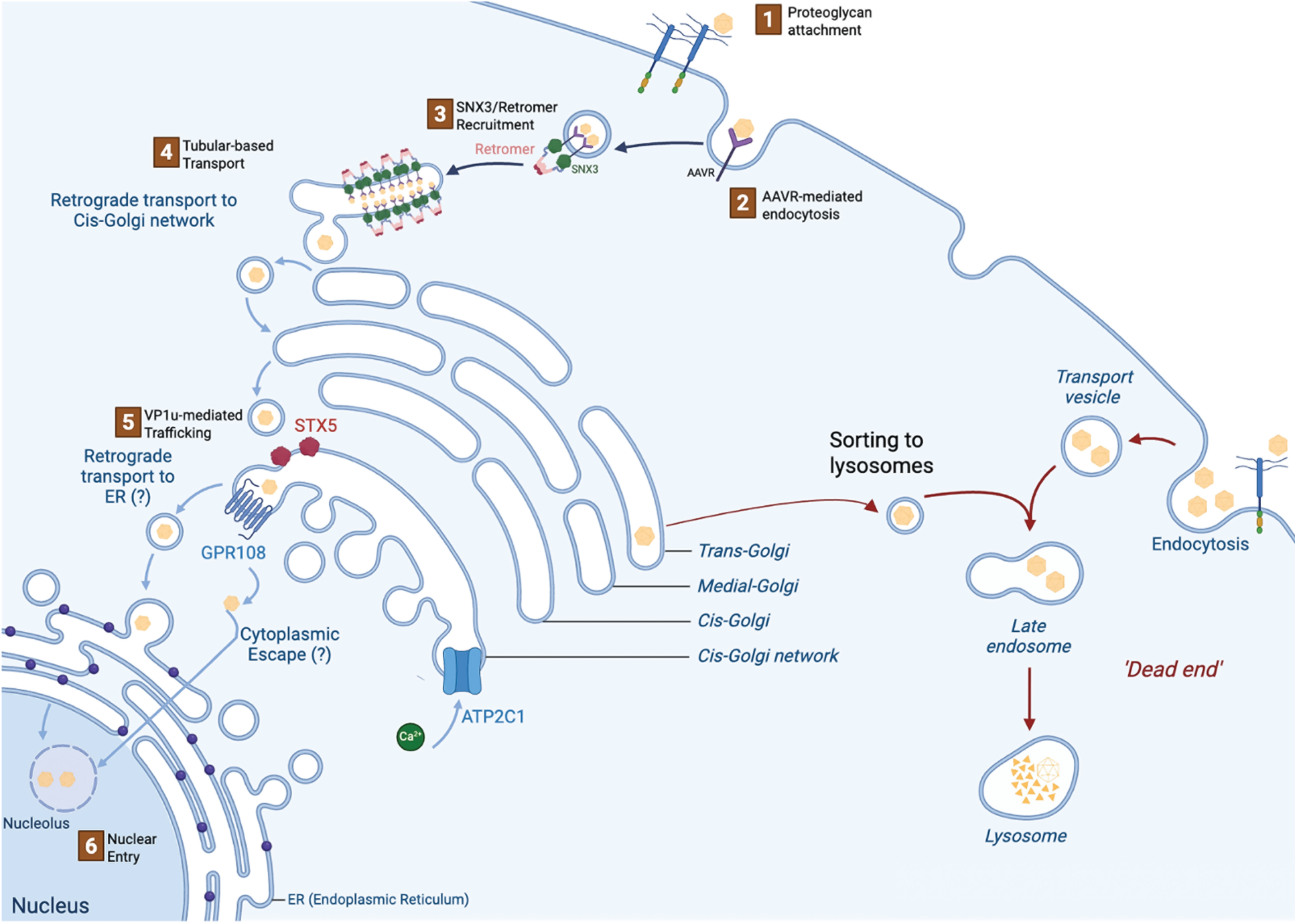
Updated model of AAV2 intracellular trafficking pathways. Schematic representation of productive and non-productive AAV2 entry routes. Following initial attachment to cell surface proteoglycans (1), AAV2 can undergo either AAVR-mediated endocytosis (2) leading to productive transduction, or AAVR-independent uptake resulting in lysosomal degradation (“dead end”). In the productive pathway, AAVR functions as a retromer cargo, recruiting the SNX3-retromer complex at endosomes (3) to facilitate tubular-based retrograde transport (4) to the cis-Golgi network. STX5, a SNARE protein at the cis-Golgi, mediates membrane fusion events during retrograde trafficking[15]. VP1u-mediated trafficking (5) becomes essential at or after the trans-Golgi network, where the Golgi-resident factor GPR108 facilitates AAV progression[37]. The calcium pump ATP2C1 at the Golgi may provide the calcium required for VP1u phospholipase A2 activity[42]. Following these Golgi-dependent steps, AAV proceeds toward the nucleus (6), though the mechanism of cytoplasmic escape remains uncertain (indicated by “?”). The ER’s role in this pathway, if any, requires further investigation. AAV particles entering through AAVR-independent pathways are sorted to lysosomes for degradation and fail to establish productive infection. This model challenges the traditional concept of direct endosomal escape, instead positioning AAV transduction as a receptor-guided, vesicle-mediated trafficking process with a critical checkpoint at the trans-Golgi network.

## Materials and Methods

### Peptides

All the synthetic peptides used for the assays were purchased from Genscript with a purity ≥95%. Peptides were dissolved in tubulation assay buffer (20 mM HEPES 7.5, 150 mM NaCl, 1 mM TCEP). See Table 1.

**Table 1.** Synthetic peptides used in this work.

| Peptide | Sequence |
| --- | --- |
| AAVR<sub>956-974</sub> | KRQKGKPKRKSKYKILDAT |
| AAVR<sub>956-967</sub> | KRQKGKPKRKSK |
| DMT1-II<sub>550-568</sub> | AQPELYLLNTMDADSLVSR |

### Lipids

All pure synthetic lipids were purchased from Avanti Polar Lipids in powdered format, including: 1,2-dioleoyl-sn-glycero-3-phosphocholine (DOPC) (catalog no. 850375P), 1,2-dioleoyl-sn-glycero-3-phosphoethanolamine (DOPE) (catalog no. 850725P), 1,2-dioleoyl-sn-glycero-3-phospho-L-serine (DOPS) (catalog no. 840035P) and 1,2-dioleoyl-sn-glycero-3-phospho-(1’-myo-inositol-3’-phosphate) (PI(3)P) (catalog no. 850150P). (see Table 2). Phospholipids (DOPC, DOPE, DOPS) were resuspended in chloroform (CHCl_3_) and phosphoinositides PI(3)P was resuspended in a mixture of CHCl_3_:MeOH:H_2_O in a 20:9:1 molar ratio.

**Table 2.** Lipids used in this work.

| Lipid | Reference | Stock solution |
| --- | --- | --- |
| 1,2-dioleoil-sn-glicero-3-fosfocolina (DOPC) | 850375P | 1mg/ml in chloroform |
| 1,2-dioleoil-sn-glicero-3-fosfoetanolamina (DOPE) | 850725P | 1mg/ml in chloroform |
| 1,2-dioleoil-sn-glicero-3-fosfo-L-serina (DOPS) | 840035P | 1mg/ml in chloroform |
| 1,2-dioleoyl-sn-glycero-3-phospho-(1'-myo-inositol-3'-phosphate) (PI(3)P) | 850150P | 1 mg/ml in chloroform:methanol:water (20:9:1) |

### Recombinant DNA Procedures

All the cDNAs used in this study encode for human proteins and were cloned into bacterial expression vectors. DNA sequence encoding full-length SNX3 was cloned into pHis-MBP-Parallel2 vector[43] to express this protein with a cleavable N-terminal 6xHis-maltose binding protein (MBP) tag. DNA encoding VPS26A was cloned using pET28-Sumo3 vector (EMBL, Heidelberg) with a cleavable 6xHis-Sumo3 tag at the N terminus. Full-length VPS35-VPS29 fusion construct was cloned into pGST-Parallel2 vector[43] with a cleavable N-terminal Glutathione S-transferase (GST) tag. All constructs were verified by DNA sequencing.

### Protein Expression

All proteins were expressed in *Escherichia coli* BL21 (DE3) strain. This strain grown in Luria-Bertani (LB) broth at 37°C and protein expression was induced with 0.5 mM isopropyl-β-_D_-thiogalactopyranoside (IPTG) at an optical density at 600 nm (OD_600_) of 0.6-0.8. Cells were harvested after 16-20 hr of growth at 18°C. All following purification steps were performed at 4°C.

### Protein purification

For the purification of SNX3, the cell pellet was lysed by sonication in buffer A [50 mM Tris-HCl 8, 300 mM NaCl, 1 mM dithiothreitol (DTT), 20 mM imidazole] supplemented with 0.5 mM phenylmethylsulfonyl fluoride (PMSF), 5 mM benzamidine, DNAses and lysozyme. After centrifugation at 5,000 g for 45 min, the soluble fraction was incubated for 2 h in batch with Ni^2+^-nitrilotriacetate (NTA) agarose resin (QIAGEN). After extensive washing of the beads with buffer A in a gravity column, protein was eluted in the same buffer supplemented with 250 mM imidazole. Tobacco etch virus (TEV) protease was added to the eluted sample and was incubated for 3 hr to remove the N-terminal 6×His-MBP tag. The mixture was dialyzed overnight in buffer B (25 mM Tris-HCl 6.5, 25 mM NaCl and 1 mM DTT) and a second Ni^2+^-NTA incubation was carried out to remove the cleaved tag. Finally, the protein was successfully purified using a combination of ion-exchange chromatography (HiTrapSP, GE Healthcare) using a linear gradient of NaCl followed by size-exclusion chromatography (Superdex 75 16/60, GE Healthcare) in buffer C (25 mM Tris-HCl 7.5, 300 mM NaCl and 1 mM DTT).

Isolation of retromer complex (VPS26-VPS35-VPS29) was carried out by mixing the cell pellets of GST-VPS35-VPS29 and His-Sumo3-VPS26A. Both cell pellets were lysed by high-pressure homogenization (25 kpsi) in buffer D (50 mM Tris-HCl 8.0, 500 mM NaCl, 1 mM DTT) supplemented with 0.5 mM PMSF, 5 mM benzamidine, DNAses and lysozyme and the lysate was cleared by centrifugation at 5,000 g for 45 min. Supernatant was incubated overnight in batch with glutathione-Sepharose beads (GE Healthcare) followed by an extensive washing of these beads with buffer D in a gravity column. Protein was released of the beads by 4 hr cleavage of the GST-tag with TEV protease in the same buffer and cleaved protein was incubated 2 hr in batch with Ni^2+^-NTA agarose resin in buffer D supplemented with 20 mM of imidazole. Protein was eluted with buffer D supplemented with 250 mM imidazole and SENP2 was added for the cleavage of the 6xHis-Sumo3-tag by overnight dialysis in buffer E (50 mM Tris-HCl 8.0, 150 mM NaCl, 20 mM imidazole, 1 mM DTT). Retromer was finally purified by ion-exchange chromatography (HiTrapQ, GE Healthcare) using a gradient of NaCl followed by size-exclusion chromatography (Superdex 200 16/60, GE Healthcare) in buffer C.

The concentration of all purified proteins was calculated using the theoretical extinction coefficient obtained from ExPASy.

### Humanized Mouse Liver

All animal care and experimental procedures were approved by the joint Children’s Medical Research Institute (CMRI) and The Children’s Hospital at Westmead Animal Care and Ethics Committee. CMRI’s established FRG[21] mouse colony was used to breed recipient animals. FRG mice were housed in individually ventilated cages with 2-(2-nitro-4-trifluoro-methylbenzoyl)-1,3-cyclohexanedione (NTBC) supplemented in drinking water (8 mg/mL). FRG mice, 6 to 8 weeks old, were engrafted with human hepatocytes (Lonza, Basel, Switzerland) as described previously. hFRG mice were placed on 10% NTBC prior to transduction with vectors and were maintained on 10% NTBC until harvest. Mouse livers were processed as described recently with no modifications[20]. Briefly, mouse livers fixed with 4% (w/v) paraformaldehyde, cryo-protected in 10%–30% (w/v) sucrose before freezing in O.C.T. (Tissue-Tek; Sakura Finetek USA, Torrance, CA, USA). Frozen liver sections (5 μm) were permeabilized in −20°C methanol, then at room temperature in 0.1% Triton X-100, and then reacted with an anti-AAVR antibody (Abcam, ab105385) at 1:100/Alexa Flour555 at 1:500, anti-GAPDH (ab215227) at 1:500 and DAPI (Invitrogen, D1306) at 0.08 ng/mL. After immunolabeling, the images were captured and analyzed on a Zeiss Axio Imager.M1 using ZEN 2 software.

### Culture and immunofluorescence analysis of HuH-7 cells

Cells were cultured in Dulbecco’s modified Eagle’s medium (DMEM) (Gibco, 11965-092) supplemented with 100 U/mL penicillin/100 μg/mL streptomycin (Sigma-Aldrich, P4458) and 10% fetal bovine serum (FBS) (Sigma-Aldrich) and non-essential amino acids (Gibco, 11140-050). Cells were passaged using TrypLE express enzyme (Gibco, 12604-21).

For transduction studies, cells were seeded on coverslips in 24-well plates in complete DMEM at a density of 1 × 10^5^ cells per well and allowed to adhere overnight in a tissue culture incubator at 37 °C and 5% CO_2_. MOI of 10,000 AAV2_ss-LSP1-eGFP-WPRE-BGHpA were added to the wells next day and coverslips were harvested at 6hrs, 16hrs, 48hrs and 72hrs post transductions. For immunofluorescence, cells were washed 1 x in PBS, then fixed in 4% paraformaldehyde, permeabilized with 0.2% Triton X-100, and blocked in 10% normal donkey serum diluted in 1× PBS. Cells were reacted with anti-AAVR antibody (Abcam, ab105385) at 1:100/Alexa Fluor 594 at 1:500, anti-giantin antibody (BioLegend, 908701)/Alexa Fluor 488 at 1:500, anti-TGN46/Alexa Fluor 488 (Abcam, ab306606) at 1:500, A20 antibody (ARP 03-65155) at 1:20/Alexa Fluor 594 at 1:500, Phallodin-iFluor 647 (Abcam ab176759) and DAPI (Invitrogen, D1306) at 0.08 ng/mL. After immunolabeling, the images were captured and analyzed on a Zeiss Axio Imager.M1 using ZEN 2 software.

### AAV vector packaging and viral production

AAV constructs were packaged into AAV capsids using HEK293 cells and a helper-virus-free system as previously described[44]. All vectors/viruses were purified using iodixanol gradient ultracentrifugation as previously described[45] AAV preparations were titered by droplet digital PCR (ddPCR) using EGFP-specific primers EGFP-forward/reverse (For/Rev) (Table 3).

**Table 3.**
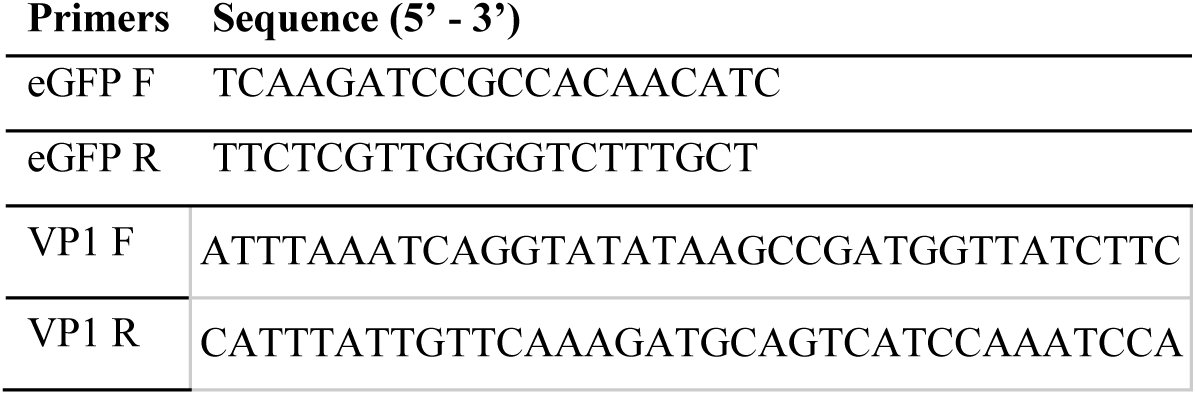
Oligonucleotides used in this work.

### HuH-7 AAVR-KO cell line

The HuH-7 AAVR-knockout cell line used in this study was generated and described previously (34095344) and has been validated for loss of AAVR expression.

### Human liver sections

Human liver samples used for AAVR immunostaining were previously described and characterized in Cabanes-Creus et al.[46]. In brief, livers unsuitable for transplantation but consented for research were obtained through DonateLife under protocols approved by the Sydney Local Health District Ethics Review Committee (X18-0523 & 2019/ETH08964). Tissue procurement, processing, and immunofluorescence staining were performed exactly as outlined in the cited study.

### HC RNA-FISH

RNA fluorescence in situ hybridization was performed using the HCR Pro RNA-ISH kit (Molecular Instruments) following the manufacturer’s protocol. A custom probe targeting Cerulean (5’-AGCTGGAGTACAACGCCATCAGCG-3’) was hybridized at 37 °C, and signal was amplified by hybridization chain reaction with fluorophore-labeled hairpins. Slides were counterstained with DAPI and mounted in antifade medium for image capture and analysis on Zeiss LSM 800-Airyscan microscope using ZEN Black software.

### AAV Variants

The described AAV2 variants throughout this manuscript were generated with site-directed mutagenesis using the overlapping primers described in Table 3. Briefly, the AAV2-VP1 mutant was generated by mutating the start ATG codon on VP1, and the PLA2-deficient variant was generated by mutating amino acids 75 and 76 (HD) to AN, as previously described[14].

### Lipids

All pure synthetic lipids were purchased from Avanti Polar Lipids in powdered format, including: 1,2-dioleoyl-sn-glycero-3-phosphocholine (DOPC) (catalog no. 850375P), 1,2-dioleoyl-sn-glycero-3-phosphoethanolamine (DOPE) (catalog no. 850725P), 1,2-dioleoyl-sn-glycero-3-phospho-L-serine (DOPS) (catalog no. 840035P) and 1,2-dioleoyl-sn-glycero-3-phospho-(1’-myo-inositol-3’-phosphate) (PI(3)P) (catalog no. 850150P). (see Table 2). Phospholipids (DOPC, DOPE, DOPS) were resuspended in chloroform (CHCl_3_) and phosphoinositides PI(3)P was resuspended in a mixture of CHCl_3_:MeOH:H_2_O in a 20:9:1 molar ratio.

### Liposome Preparation

Liposomes for tubulation assays were produced by mixing DOPC, DOPE, DOPS and PI(3)P in a 45:30:20:5 molar ratio respectively. For proper mixture homogenization, lipids were incubated 1 h at 37°C. Mixed lipids were extensively dried in a vacuum desiccator and re-hydrated in tubulation assay buffer (20 mM HEPES 7.5, 150 mM NaCl, 1 mM TCEP) at a final lipid concentration of 1.5 mM. Resuspended mixture was heated 1 h at 60°C to facilitate lipid solubilization in the buffer. This suspension of multilamellar liposomes was subjected to 11 cycles of freeze-thawed-vortex (in liquid nitrogen and 50°C waterbath, respectively) to produce unilamellar liposomes and extruded through a 0.2 μM pore-size polycarbonate filter by a mini-extruder (Avanti Polar Lipids) 11 times. Liposomes were stored at 4°C under argon atmosphere and used for up to three days.

### Tubulation Assays

For tubulation reactions, 2.5 μM of the retromer complex, 8.75 μM of SNX3 and 25 μM of AAVR_956-974_ or its mutants or DMT1-II_550-568_ were incubated with 600 μM of liposomes overnight at 4°C in tubulation assay buffer. Four microliters of the sample were frontside-blotted for 2 s at a relative humidity of 90% and a temperature of 8°C on a glow-discharged holey carbon grid (Quantifoil® R2/2 Cu 200 mesh grids) before plunge-freezing in liquid ethane (Leica Plunger EM P2). Images were adquired on a JEM-2200FS electron microscope operated at 200 kV using a K2 direct detector.

### Structural modelling

The amino acid sequences of human AAVR (residues 956–974; UniProt Q8IZA0), VPS35 (residues 1–470; UniProt Q96QK1), VPS26A (residues 1–327; UniProt O75436), and SNX3 (residues 1–162; UniProt O60493) were uploaded as individual entities to AlphaFold3 for complex structure prediction using the AlphaFold Server (https://alphafoldserver.com) with default parameters. Five independent models of the AAVR peptide–retromer–SNX3 complex were generated with different random seeds. Model quality, assessed using AlphaFold3-derived pLDDT and PAE scores, showed minimal variation among predictions. Model visualization and figure generation were performed in PyMOL.

### Endosomal escape assay

HepG2 cells were genetically modified to stably express eGFP-Galectin 8 (Gal-8) and were used for endosomal escape assays (produced by Genscript). eGFP puncta counting was used as a proxy for endosomal escape as reported elsewhere[31]. In brief Gal-8 HepG2 cells were plated on rat tail collagen (08-115, Sigma) coated flat-bottom plates at a density of 20,000 cells per well 24 hours before AAV/LNP treatment. The cells were treated with 1 μg/mL recombinant ApoE4 (350-04, Peprotech, Cranbury, NJ) and the desired concentration of LNP and AAV simultaneously. To monitor productive delivery of payload following endosomal escape AAV2 was used with NLS-mScarlett payload while LNPs encapsulated mCherry mRNA (obtained from Trilink). Live cell imaging was conducted with an Opera Phenix Spinning Disc Confocal set in epifluorescence mode using a 40x water objective. A Harmony software algorithm for counting puncta within cell boundaries were utilized for eGFP Gal-8 puncta quantification over time. Similarly, a Harmony algorithm for quantifying mCherry and mScarrlett positive cells vs total eGFP Gal-8 HepG2 populations was employed to track productive delivery of payload.

### LNP formulation and characterization

LNPs were prepared using SM-102 (33474, Cayman Chemical, Ann Arbor, MI), 1,2-distearoyl-sn-glycero-3-phosphocholine (DSPC), cholesterol, and C14-polyethylene glycol (PEG) (Avanti Polar Lipids, Alabaster, AL) at a molar ratio of 50:10:38.5:1.5. The lipids were dissolved in ethanol and combined with 50 mM, pH 3 citrate buffer containing nucleic acid payload at a flowrate ratio of 3:1 (aqueous:ethanol) using a microfluidic device (Precision Nanosystems, Vancouver, Canada). The total amine:phosphate (N:P) ratio between ionizable lipids and nucleic acids was fixed at 4.92. After synthesis, formulations were purified in dialysis cassettes (Slide-A-Lyzer, 20 KDa molecular weight cutoff [MWCO], Thermo Fisher Scientific, Waltham, MA) with phosphate-buffered saline (PBS) (pH 7.4, no Ca^2+^ or Mg^2+^) overnight for the removal of excess ethanol. Formulations were then concentrated using Amicon filters (20 kDa MWCO; Sigma-Aldrich, St Louis, MO) and passed through syringe filters (Acrodisc 0.45µm PTFE 13mm, Cytiva). We added 10% sucrose (prepared from powdered sucrose stock; Sigma-Aldrich) to the formulations prior to storage at −80°C.

## Funding

This research was supported by the Spanish State Research Agency through Project PID2023-151986NB-I00 (to A.H.).

## Competing interests

M.P.P., A.L.R., A.H., declare that they have no competing interests.

## Notes

### Competing Interest Statement

The authors have declared no competing interest.

