## Supplementary Figures for "AAV2 Bypasses Direct Endosomal Escape by Using AAVR to Access the Trans-Golgi Network en Route to the Nucleus"

**Supplementary Figure 1.**

Transmission electron microscopy (TEM) of liposomes incubated with individual or pairwise components. Neither retromer, SNX3, nor the AAVR cytosolic tail peptide (residues 956–974) alone, nor any two-component combination, is sufficient to induce tubulation. Robust tubule formation requires the concerted presence of retromer, SNX3, and a cargo peptide.

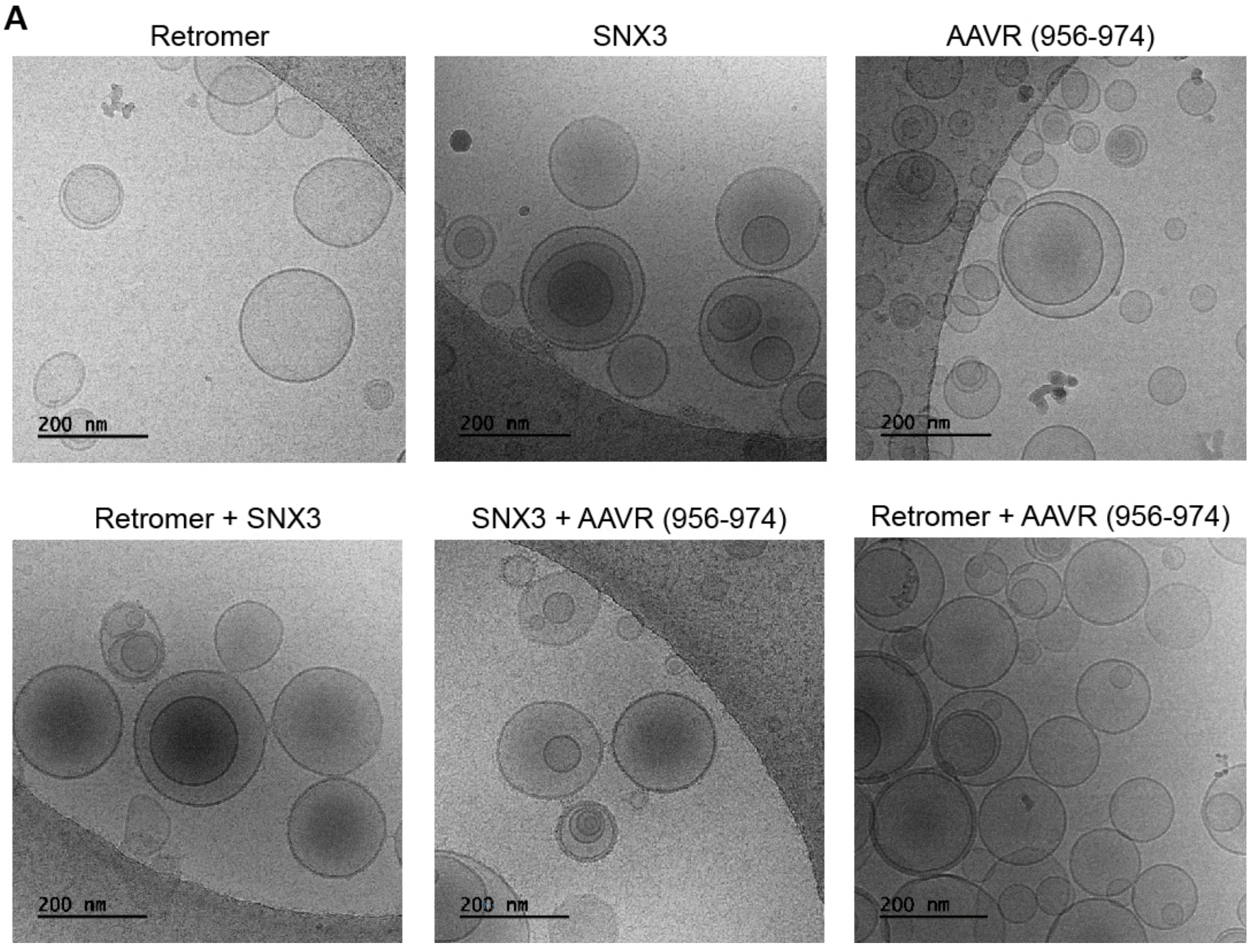
